# Assessing Bacterial contaminants on hospital devices at Zomba Central Hospital Intensive care Unit (ICU) and Theatre

**DOI:** 10.64898/2026.09.14.751631

**Authors:** Nelson Chimwaza, David Kulapani, Tonney Nyirenda

## Abstract

Hospital devices in intensive care unit (ICU) and theatre settings are prone to bacterial contamination, posing a major risk to patient safety. No previous study has been conducted at Zomba Central Hospital (ZCH) to assess such contamination. This study aimed to determine the prevalence and types of bacteria contaminating ICU and theatre devices, evaluate the effectiveness of cleaning protocols, and assess healthcare workers’ knowledge on infection control. A cross-sectional study involving 25 hospital devices and 30 healthcare workers was conducted. A self-administered questionnaire assessed staff knowledge and perceptions, while swabs were collected from devices before and after cleaning for culture and antimicrobial susceptibility testing. Of the 25 swabs collected, 72.0% were contaminated before cleaning, decreasing significantly to 36.0% after cleaning (p = 0.049). Gram-positive bacteria predominated (69.0%), with ventilators being the most contaminated devices. Multidrug resistance was detected in extended-spectrum beta-lactamase (ESBL)–producing *Klebsiella pneumoniae, Enterobacter* spp., and carbapenem-resistant *Serratia marcescens*. Although most staff acknowledged the importance of infection prevention (93.3%), knowledge of bacterial contaminants (73.3% low) and recommended cleaning methods (90.0% low) was poor. Lower cadre roles (OR = 0.30, p = 0.020) and lower education levels (OR = 4.87, p = 0.032) were significantly associated with reduced knowledge. The study concludes that ICU and theatre devices at ZCH are contaminated with diverse bacterial species, including multidrug-resistant strains. While existing cleaning protocols significantly reduce contamination, enhanced staff training and standardized cleaning procedures are essential to ensure aseptic hospital environments.

## 1. Introduction

Hospital devices in intensive care unit (ICU) and theatre settings are prone to bacterial contamination, posing a significant risk to patient safety. Contaminated devices can serve as reservoirs for healthcare-associated pathogens and facilitate cross-transmission between patients and healthcare workers, particularly in high-risk environments such as ICUs and operating theatres.

Studies have consistently demonstrated high rates of bacterial contamination on hospital equipment. For example, Bhatta et al. (2022) reported contamination of 64.7% of ICU devices in Nepal, while Mutib et al. (2021) found that 33.3% of theatre devices were contaminated prior to fumigation in Iraq. Even higher contamination rates have been documented in Ethiopia, where 88.5% of swabs from medical equipment and inanimate surfaces yielded bacterial growth (Darge et al., 2019). Such findings highlight the persistence of bacterial contamination despite routine cleaning practices.

Commonly isolated organisms from hospital devices include *Staphylococcus aureus, Pseudomonas aeruginosa*, and *Acinetobacter baumannii*, all of which are associated with severe healthcare-associated infections (Weber et al., 2017). Device-associated infections contribute substantially to morbidity, mortality, and healthcare costs, particularly when multidrug-resistant organisms are involved (Brito, 2017; Percival et al., 2015).

In Malawi, healthcare-associated infections remain a major public health concern. Point prevalence surveys at Queen Elizabeth Central Hospital have reported HAI prevalence ranging from 9.6% to 11.4% (Bunduki et al., 2021; Kachipedzu et al., 2024). Despite this burden, there are no local data describing bacterial contamination of hospital devices at Zomba Central Hospital. Generating such evidence is essential to inform infection prevention and control (IPC) strategies and antimicrobial stewardship interventions.

This study therefore aimed to determine the prevalence and types of bacterial contaminants on ICU and theatre devices at ZCH, evaluate the effectiveness of current cleaning and disinfection protocols, assess antimicrobial susceptibility patterns of isolated bacteria, and examine healthcare workers’ knowledge regarding device contamination and infection control.

## 2. Materials and Methods

### 2.1 Study Design and Setting

A cross-sectional study was conducted at Zomba Central Hospital, one of the four tertiary referral hospitals in Malawi. The hospital serves as a referral Centre for several district and mission hospitals in the eastern region and has a six-bed ICU and a high-volume operating theatre performing approximately 15 procedures per day.

### 2.2 Study Population and Sampling

The study included reusable hospital devices that come into direct contact with patients in the ICU and theatre, as well as healthcare workers assigned to these units. Twenty-five (25) hospital devices and thirty (30) staff members were selected using stratified sampling based on department (ICU and theatre) and device type or job role, respectively.

### 2.3 Data Collection

#### 2.3.1 Device Swabbing and Microbiological Analysis

Swabs were collected from selected devices before and after routine cleaning using sterile cotton swabs moistened with sterile normal saline. An approximate surface area of 5 cm × 5 cm was swabbed for each device. Post-cleaning swabs were taken from the same site 10–15 minutes after cleaning. Samples were transported to the microbiology laboratory within 30 minutes.

Swabs were cultured on Blood Agar and MacConkey Agar and incubated at 37°C for 48 hours. Bacterial isolates were identified using standard microbiological techniques, including Gram staining and biochemical tests. Antimicrobial susceptibility testing was performed using the disk diffusion method according to ZCH laboratory standard operating procedures.

#### 2.3.2 Staff Knowledge Assessment

A structured, self-administered questionnaire was used to assess staff knowledge and perceptions regarding bacterial contamination and infection control practices. The tool was developed following literature review, expert consultation, pilot testing, and validation procedures.

### 2.4 Data Analysis

Data was entered into Microsoft Excel and analyzed using SPSS version 20. Descriptive statistics were used to summarize contamination prevalence and staff knowledge levels. Chi-square tests assessed the effectiveness of cleaning protocols, while ordinal logistic regression identified predictors of staff knowledge.

### 2.5 Ethical Considerations

Ethical approval was obtained from the College of Medicine Research Ethics Committee (COMREC; P.10/24-1163). Permission to conduct the study was granted by ZCH management. Written informed consent was obtained from all participants, and confidentiality was maintained throughout the study.

## 3. Results

### 3.1 Prevalence and Types of Bacterial Contaminants

Overall, 72.0% (18/25) of devices were contaminated before cleaning, compared to 36.0% (9/25) after cleaning (Figure 1). Pre-cleaning contamination was 73.3% in ICU devices (Table 1) and 70.0% in theatre devices (Table 2). Gram-positive bacteria accounted for 69.0% of isolates, while Gram-negative bacteria constituted 31.0% (Figure 2).

**Figure 1:**
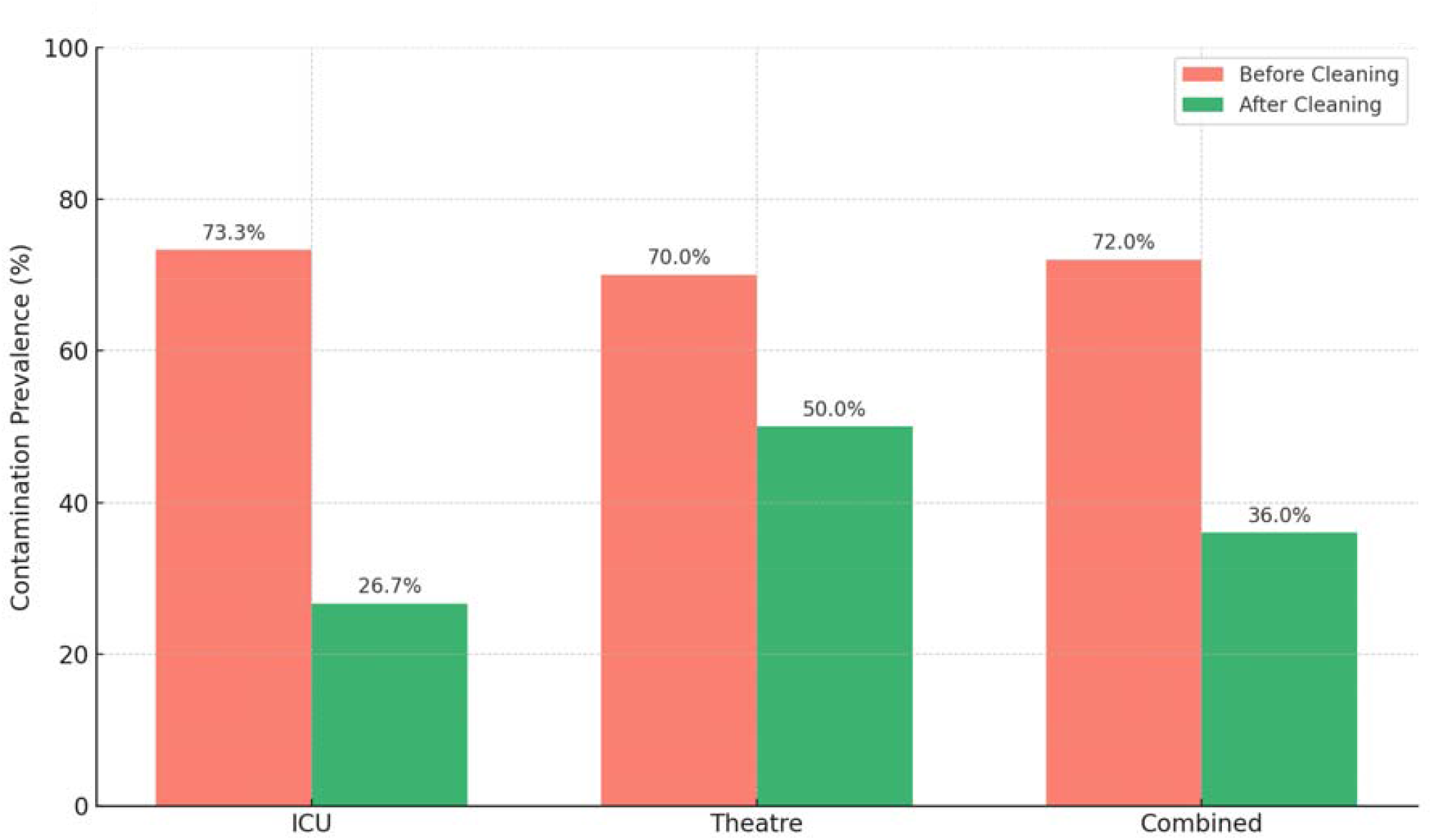
Contamination prevalence in ICU and Theatre before and after cleaning.

**Table 1.**
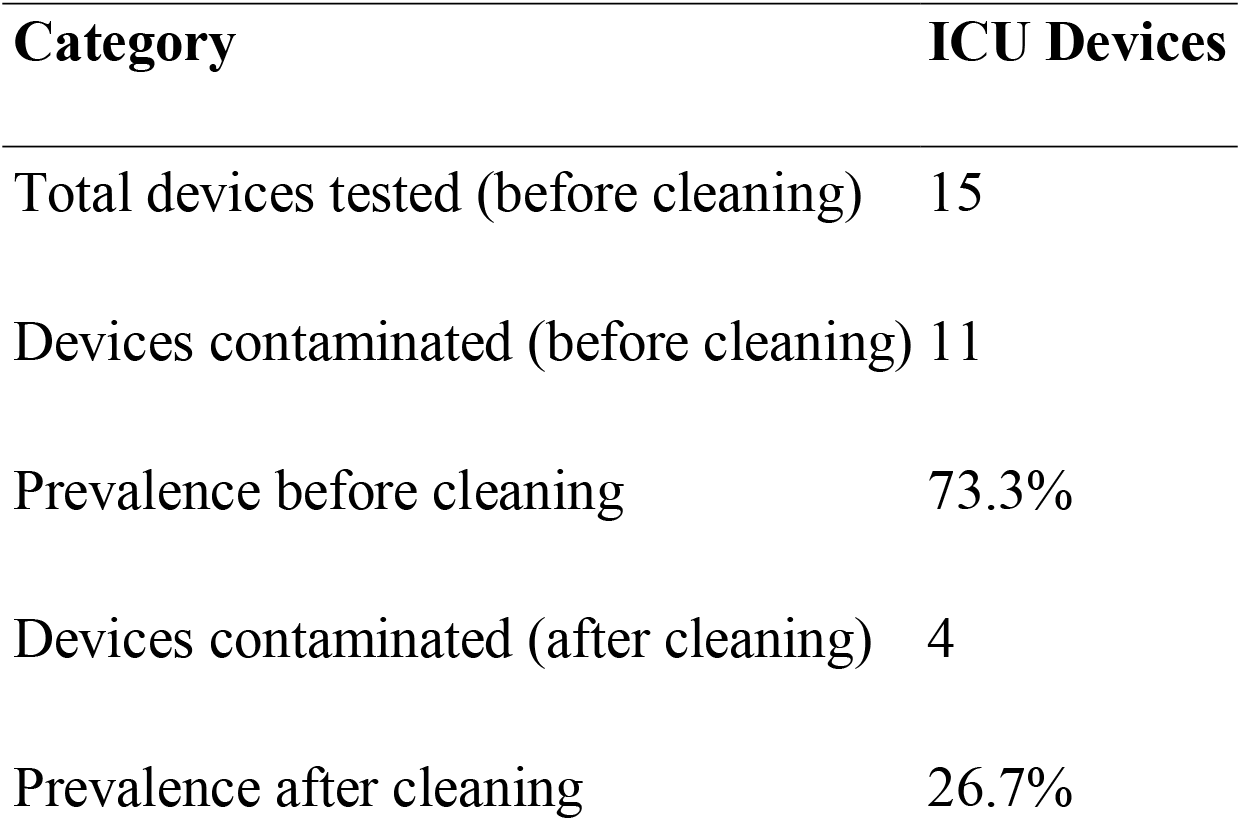
Prevalence of bacterial contaminants on hospital devices at ICU.

| Category | ICU Devices |
| --- | --- |
| Total devices tested (before cleaning) | 15 |
| Devices contaminated (before cleaning) | 11 |
| Prevalence before cleaning | 73.3% |
| Devices contaminated (after cleaning) | 4 |
| Prevalence after cleaning | 26.7% |

**Table 2.** Prevalence of bacterial contaminants on hospital devices at Theatre.

| Category | Theatre Devices |
| --- | --- |
| Total devices tested (before cleaning) | 10 |
| Devices contaminated (before cleaning) | 7 |
| Prevalence before cleaning | 70.0% |
| Devices contaminated (after cleaning) | 5 |
| Prevalence after cleaning | 50.0% |

**Figure 2:**
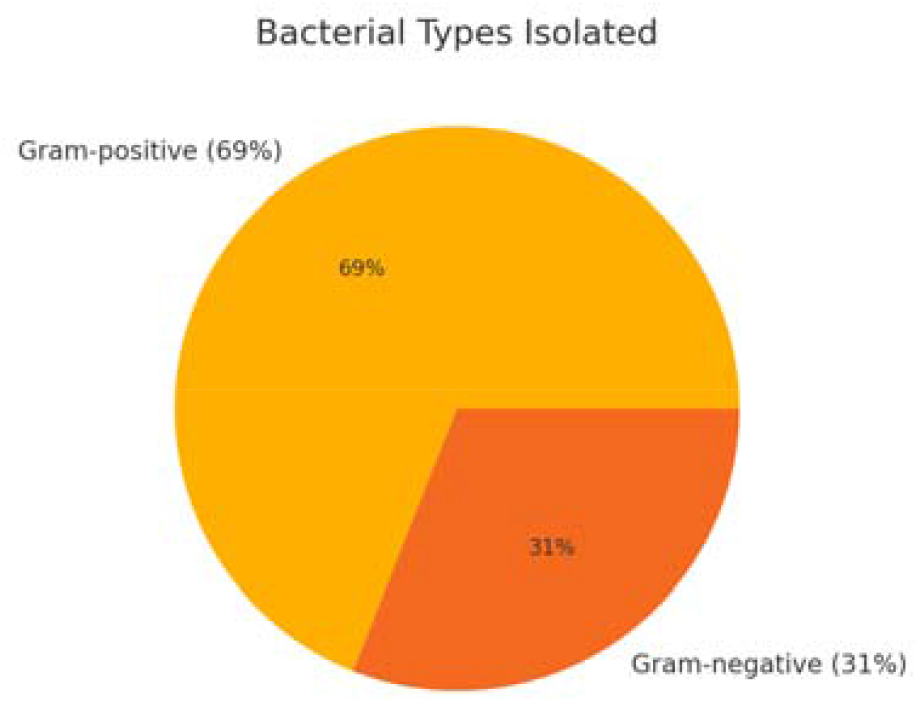
Distribution of gram-positive vs. gram-negative bacteria.

Ventilators were the most contaminated devices in both settings. Identified organisms included methicillin-resistant coagulase-negative staphylococci (MRCoNS), *Bacillus* spp., *Micrococcus* spp., *Pseudomonas aeruginosa*, ESBL-producing *Klebsiella pneumoniae* and *Enterobacter* spp., and carbapenem-resistant *Serratia marcescens* (Table 3 and Figure 3).

**Table 3.**
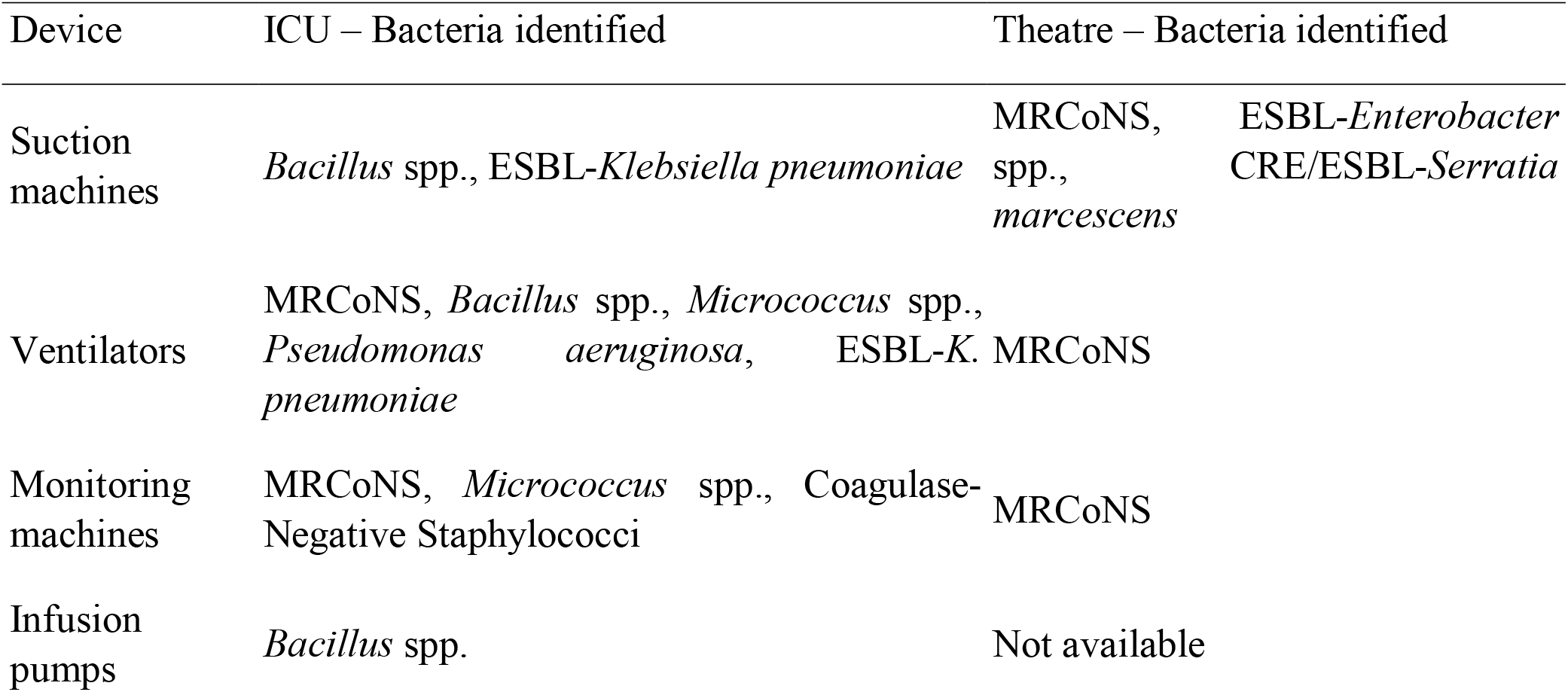
Summary of device-specific bacteria in ICU and Theatre.

**Figure 3:**
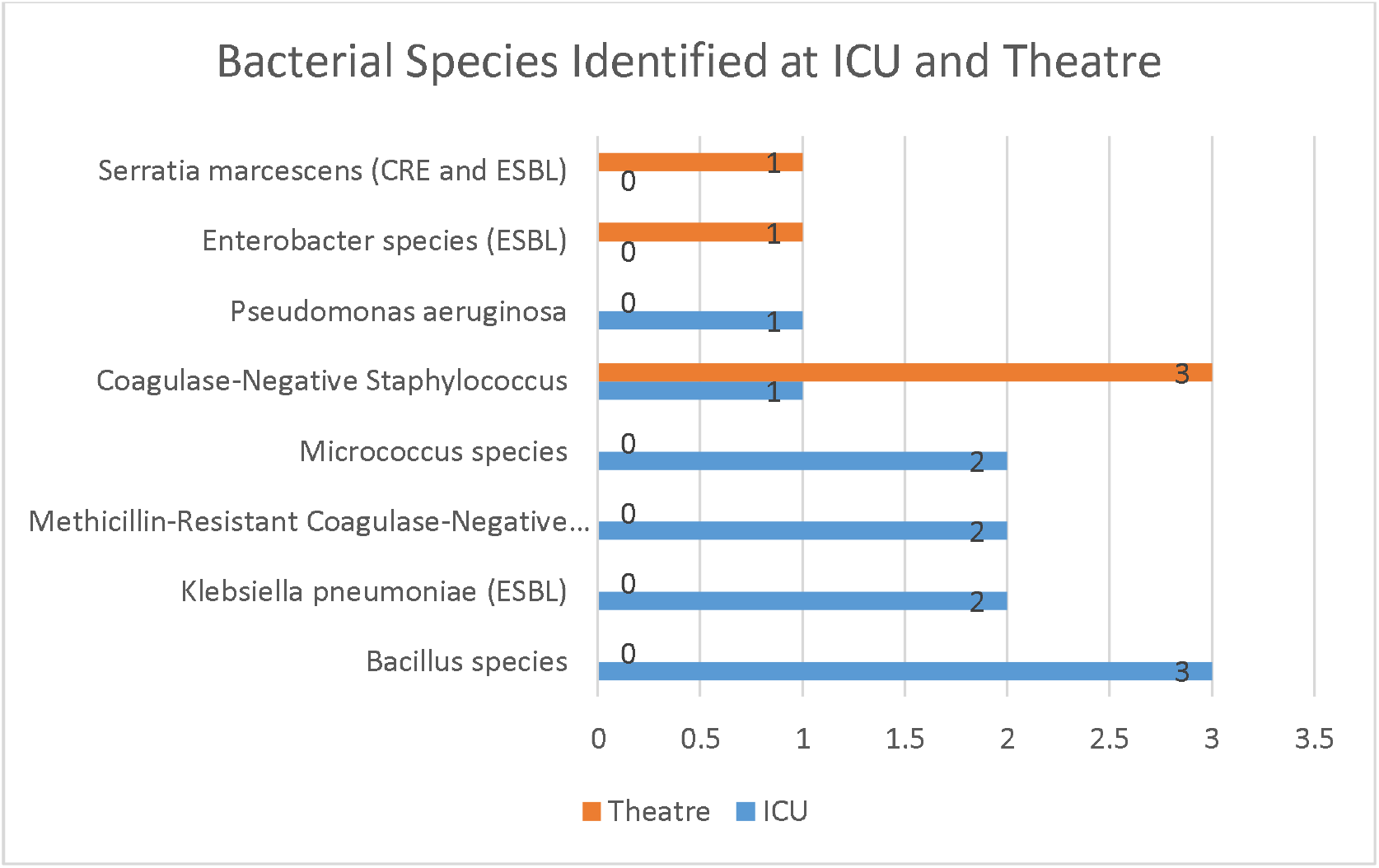
Overall number of bacteria species identified at ICU and Theatre.

### 3.2 Effectiveness of Cleaning and Disinfection

A significant reduction in bacterial contamination was observed after cleaning (χ^2^ = 7.86, p = 0.049), indicating that current cleaning and disinfection protocols were effective, although residual contamination persisted, particularly in theatre devices.

### 3.3 Antimicrobial Susceptibility Patterns

High levels of antimicrobial resistance were observed. Gram-positive isolates showed marked resistance to penicillin and erythromycin, while Gram-negative isolates demonstrated resistance to ampicillin, ceftriaxone, and Ceftazidime. ESBL and carbapenem resistance were detected among enterobacterales (Table 4 and Figure 4).

**Table 4.** Antimicrobial susceptibility patterns of isolated bacteria.

| Bacteria isolated | Sensitive to | Resistant to |
| --- | --- | --- |
| *Bacillus species* | Meropenem, Vancomycin, Erythromycin | Ciprofloxacin |
| *Klebsiella pneumoniae* (ESBL) | Amikacin, Tigecycline, Meropenem | Ampicillin, Ceftazidime, Ceftriaxone, Ciprofloxacin, Cefepime, Gentamicin |
| *Methicillin-Resistant Coagulase-Negative Staphylococcus* (MRCoNS) | Tigecycline, Amikacin, Gentamicin | Penicillin, Cefoxitin, Ciprofloxacin, Tetracycline, Erythromycin |
| *Micrococcus species* | Gentamicin, Tigecycline, Cefoxitin, Amikacin | Erythromycin, Penicillin, Ciprofloxacin |
| *Pseudomonas aeruginosa* | Meropenem, Amikacin | Ceftazidime, Cefepime, Ciprofloxacin |
| *Enterobacter species* (ESBL) | Meropenem, Ciprofloxacin, Amikacin | Ceftriaxone, Ceftazidime, Ampicillin |
| *Serratia marcescens* (CRE and ESBL) | Tigecycline | Meropenem, Ciprofloxacin, Ceftriaxone, Gentamicin, Ceftazidime, Ertapenem |

**Figure 4.**
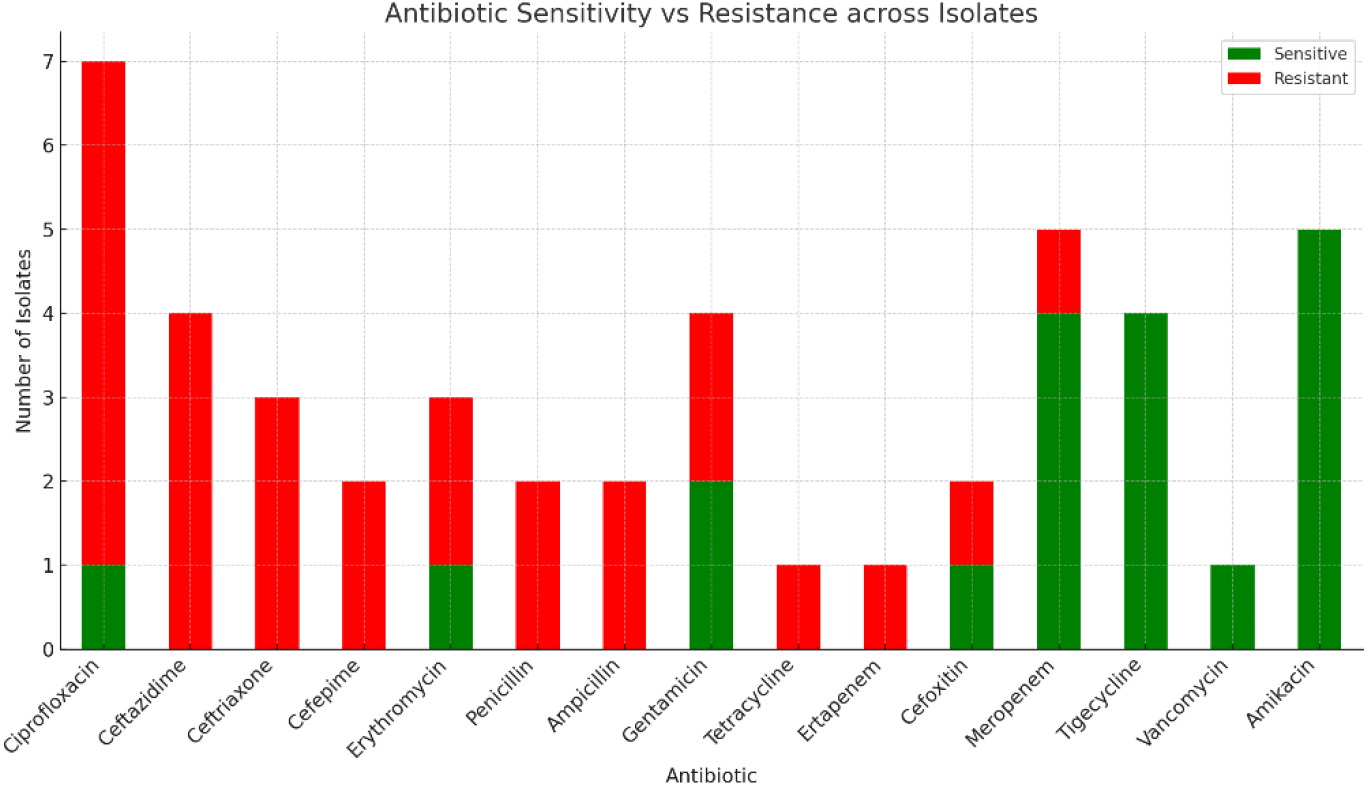
Antibiotic sensitivity vs. resistance across isolates.

### 3.4 Staff Knowledge on Infection Control

Only 6.7% of staff demonstrated high knowledge regarding bacterial contaminants, while 73.3% had low knowledge (Table 5). Ordinal logistic regression showed that lower cadre occupation (OR = 0.30, p = 0.020) and lower education level (OR = 4.87, p = 0.032) were significantly associated with lower knowledge levels (Figure 5).

**Table 5:** Ordinal regression of predictors of knowledge level among staff.

| Predictor | Estimate (B) | Std. Error | p-value | Odds Ratio (e<sup>B</sup>) | 95% CI (Lower–Upper) |
| --- | --- | --- | --- | --- | --- |
| Occupation | -1.200 | 0.516 | 0.020 | 0.30 | 0.11 – 0.83 |
| Education Level | 1.584 | 0.737 | 0.032 | 4.87 | 1.15 – 20.68 |

**Figure 5:**
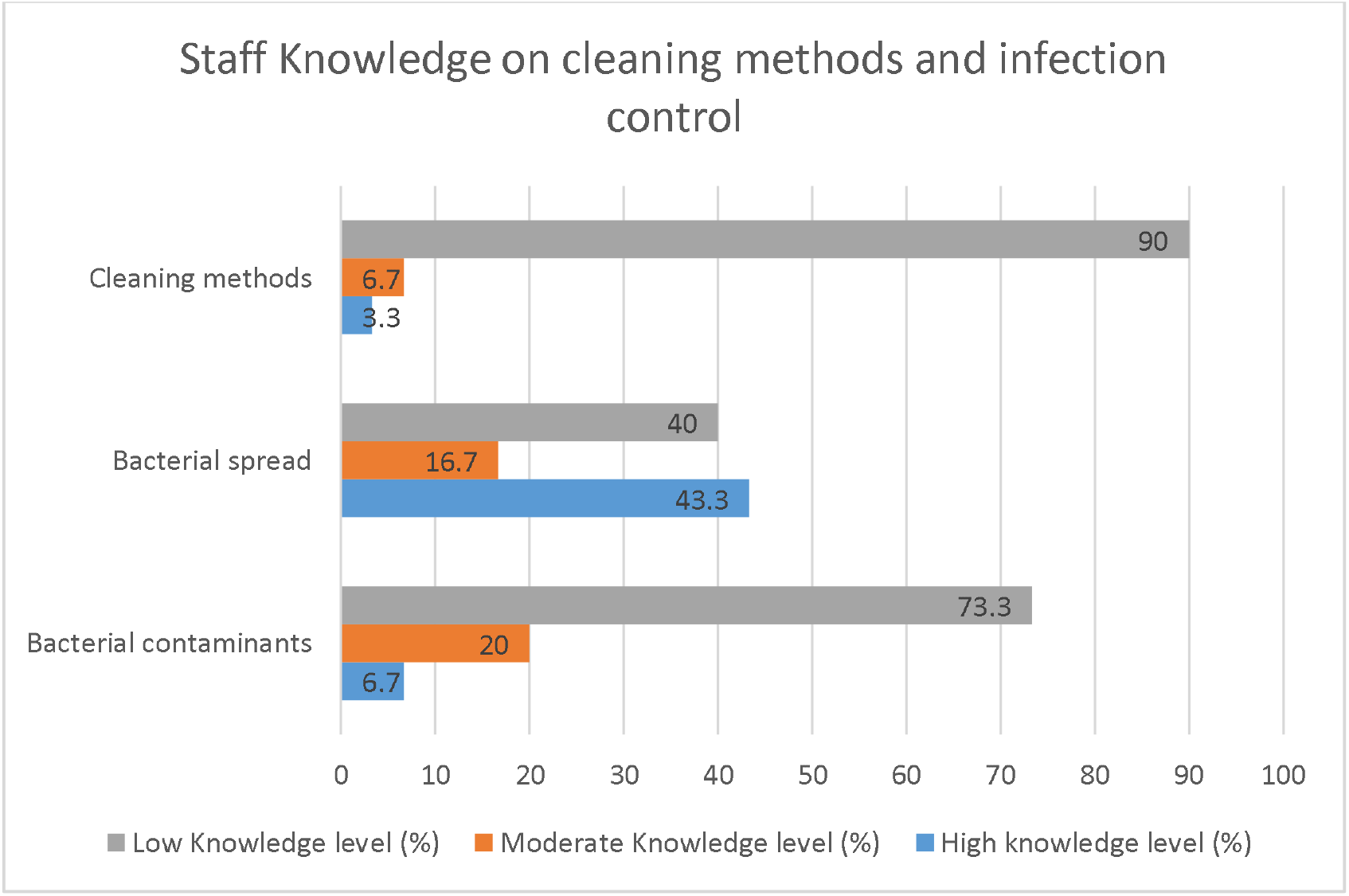
Staff knowledge on cleaning methods and infection control.

## 4. Discussion

This study demonstrated a high level of bacterial contamination on ICU and theatre devices at ZCH, consistent with findings from similar resource-limited settings (Darge et al., 2019; Sebre et al., 2020). The predominance of Gram-positive organisms, particularly MRCoNS, aligns with reports from Nigeria and Ethiopia (Ndu et al., 2022; Sebre et al., 2020).

Although current cleaning protocols significantly reduced contamination, residual contamination especially in theatre devices suggests the need for improved and standardized cleaning procedures. The detection of ESBL and carbapenem-resistant organisms is of particular concern and underscores the importance of strengthening antimicrobial stewardship and IPC interventions.

The low level of staff knowledge regarding recommended cleaning methods highlights a critical gap in infection control training. Similar associations between staff cadre, education level, and IPC knowledge have been reported elsewhere (Russotto et al., 2015; Erasmus et al., 2015)

### 4.1 Limitations

The study included fewer devices than initially planned due to limited availability. In addition, not all antimicrobial agents currently in use were tested, and the effectiveness of different disinfectants was not compared.

### 4.2 Conclusions

Hospital devices in the ICU and theatre at ZCH are contaminated with diverse bacterial species, including multidrug-resistant organisms. While existing cleaning protocols significantly reduce bacterial load, contamination persists, particularly in theatre settings. Strengthening standardized cleaning procedures, regular infection control training, and antimicrobial stewardship programs is essential to improve patient safety.

## 5. Acknowledgements

The author would like to thank and praise God for his blessings and grace. Second, I would like to extend my gratitude to Dr David Kulapani and Associate Professor Tonney Nyirenda for their scientific knowledge and continuous support in this study.

## 6. Author’s Contribution

Nelson Chimwaza: Conceptualization, Data Pre-processing, Data analysis, Manuscript preparation David Kulapani: Reviewing and Editing, Tonney Nyirenda: Reviewing and Editing

